# Suprachoroidal Delivery of Anti-Angiogenic Peptide Microparticles Enables Sustained Activity with Favorable Ocular Safety

**DOI:** 10.64898/2026.06.30.735614

**Authors:** Adam C. Mirando, Raquel Lima e Silva, Jikui Shen, Thomas J. Robinson, Jordan J. Green, Peter A. Campochiaro, Aleksander S. Popel, Niranjan B. Pandey

## Abstract

Retinal and choroidal vascular diseases are major causes of vision loss that require frequent intravitreal anti-VEGF therapy. Anti-angiogenic peptide AXT107 demonstrated efficacy in preclinical studies and was advanced to the clinical stage. To provide for sustained delivery of the peptide and avoid complications with intravitreal injection, we evaluated suprachoroidal delivery of AXT107 microparticles (MP-AXT107). The original, soluble AXT107 formulation was ineffective at inhibiting laser-induced choroidal neovascularization (CNV) in our rat model and was consequently reformulated as microparticles. MP-AXT107 demonstrated high peptide incorporation efficiency, reproducible morphology, and physical and chemical stability for at least 9 months under refrigerated storage. In the rat CNV model, suprachoroidal MP-AXT107 significantly reduced neovascular area by approximately 60% relative to vehicle controls. Safety and durability were evaluated in a 9-month GLP toxicology study in Göttingen minipigs following a single suprachoroidal injection of vehicle or MP-AXT107 (0.125–1.25 mg/eye). Transient increases in IOP and mild ocular inflammatory findings were observed immediately following administration but resolved rapidly without lasting effects. No treatment-related adverse ocular findings were observed during the remainder of the study, and the highest tested dose (1.25 mg/eye) was established as the no-observed-adverse-effect level. Bioanalysis at study completion demonstrated persistent AXT107 localization primarily within choroid/RPE and scleral tissues, with no signs of systemic exposure. Collectively, these findings demonstrate that suprachoroidal delivery of MP-AXT107 enables sustained anti-angiogenic activity with favorable ocular safety and prolonged tissue retention, supporting further clinical development as a durable therapy for retinal and choroidal vascular diseases.

## Introduction

Retinal and choroidal vascular diseases, including neovascular age-related macular degeneration (nAMD), diabetic macular edema (DME), and proliferative diabetic retinopathy (DR), are highly prevalent causes of vision loss and constitute a major global health burden^1,2^. Despite differences in their underlying pathogenesis, treatment strategies for these conditions primarily aim to control pathological retinal and choroidal neovascularization (CNV) and the associated vascular exudation^3,4^. Elevated intraocular levels of the pro-angiogenic factor vascular endothelial growth factor A (VEGFA) are a central driver of disease progression and represent the primary therapeutic target^5,6^.

Although repeated intravitreal injections of anti-VEGFA agents have demonstrated substantial clinical benefit, real-world outcomes are frequently suboptimal ^7,8^.

Laboratory studies and clinical trials have identified additional mediators beyond VEGFA that contribute to retinal and choroidal NV and exudation, including other VEGF family members such as VEGF-C and VEGF-D (ClinicalTrials.gov Identifier: NCT03345082), platelet-derived growth factor B (PDGFB), hepatocyte growth factor (HGF), and the Tie2 antagonists angiopoietin-2 (Ang2) and vascular endothelial cell protein tyrosine phosphatase (VE-PTP)^9–14^. Moreover, the chronic nature of these diseases and the irreversible damage associated with delayed or inadequate treatment necessitate adherence to regular, lifelong intravitreal injections to preserve vision, which in many cases poses a barrier to patient compliance. Consequently, there is a clear need for more durable therapies that target additional pathogenic signaling pathways beyond VEGFA^15,16^.

AXT107 is a 20-mer anti-angiogenic peptide with several promising characteristics for the treatment of retinal vascular diseases. Rather than directly targeting growth factor receptors, AXT107 modulates their function by binding the integrin co-factors αvβ3 and α5β1^17,18^. Through this mechanism, AXT107 inhibits the pro-angiogenic receptors VEGFR2, PDGFRβ, and cMet, while simultaneously potentiating activation of the vascular-stabilizing receptor Tie2 by its normally inhibitory ligand Ang2^19–22^. Activation of Tie2 was shown to reduce vascular permeability by stabilizing cell–cell junctions and inhibiting TNFα-mediated inflammation^22,23^. AXT107 also exhibited remarkable durability by self-assembling into a depot within the vitreous of animal models that continuously released efficacious amounts of peptide for more than 2 months^21^. Following a successful toxicology program in both rabbits and minipigs, AXT107 advanced to phase 1A clinical trials (NCT04697758 and NCT04746963).

In clinical studies in patients with nAMD, intravitreal injection of AXT107 resulted in gel formation that slowly eroded, providing prolonged suppression of exudation in some patients, thereby validating both αvβ3 and α5β1 as therapeutic targets and AXT107 as a viable treatment strategy. However, despite excellent safety in preclinical animal models, gel fragmentation and dispersion of fragments into the anterior chamber were observed in some patients and were associated with increased intraocular pressure (IOP) (NCT04697758 and NCT04746963). A similar phenomenon was observed in a separate clinical study in which biodegradable polymeric microparticles releasing the multikinase inhibitor sunitinib demonstrated excellent safety and prolonged efficacy (6 months) in animal models^24^, as well as prolonged suppression of exudation in patients with nAMD^25^; however, in a subset of patients, dispersion of microparticles into the anterior chamber resulted in elevated IOP (NCT03953079).

The absence of detectable material in the anterior chamber during preclinical toxicology studies suggests that standard animal models may fail to capture this clinically relevant effect. This limitation was further highlighted by an ex vivo study demonstrating accumulation of microbeads in the anterior chamber of primate eyes, but not porcine eyes^26^. To address this challenge, we investigated delivery to the suprachoroidal space (SCS) as an alternative to intravitreal administration. Injection into the SCS enables localization of the injectate near the posterior retina while physically segregating it from the anterior chamber, potentially reducing the risk of anterior segment exposure ^27^. The clinical feasibility of suprachoroidal drug administration has been established by the FDA approval of Xipere® (triamcinolone acetonide injectable suspension), which is delivered via suprachoroidal injection for the treatment of uveitic macular edema ^28^. In addition, recent advances in microneedle-based delivery systems have greatly simplified and improved the reliability of SCS administration, making this approach practical and reproducible without the need for specialized surgical training ^27,29^.

Based on these considerations, we hypothesized that suprachoroidal administration of AXT107 could maintain therapeutic exposure to the posterior segment while reducing the risk of anterior chamber migration associated with intravitreal depot formation. To test this hypothesis, we evaluated the ocular distribution, pharmacokinetics, safety, and anti-angiogenic activity of suprachoroidally administered AXT107 microparticles in preclinical models. Our findings support suprachoroidal delivery as a safe clinically translatable strategy for achieving sustained AXT107 exposure.

## Results

### Efficacy of suprachoroidal AXT107 solution

The efficacy of AXT107 administered to the SCS was first assessed in a rat model. AXT107 was prepared as it was in clinical trials (NCT04746963); that is, the lyophilized peptide was resuspended in 5% sucrose to generate an opalescent, homogeneous, solution-like dispersion (herein referred to as AXT107 solution). This AXT107 solution will self-assemble into a gel-like depot when introduced into ionic buffers, including biologicals like vitreous humor, as demonstrated in Figure 1A by depot formation in a vitreous mimetic. The efficacy of AXT107 solution delivered to the SCS was investigated using a rat laser-induced CNV model and compared to vehicle (5% sucrose). Neovascularization was induced 1 week after injection by photocoagulation and the eyes collected after another week. Analysis of retinal flat mounts revealed no apparent differences in the extent of neovascularization between the vehicle and AXT107 treated animals (Fig. 1B). Quantification of the total CNV area per eye supported this observation (Fig. 1C). Based on these data, we hypothesized that the concentrations of peptide transported across the choroid to the target neovascularization within the retina were insufficient.

**Figure 1.**
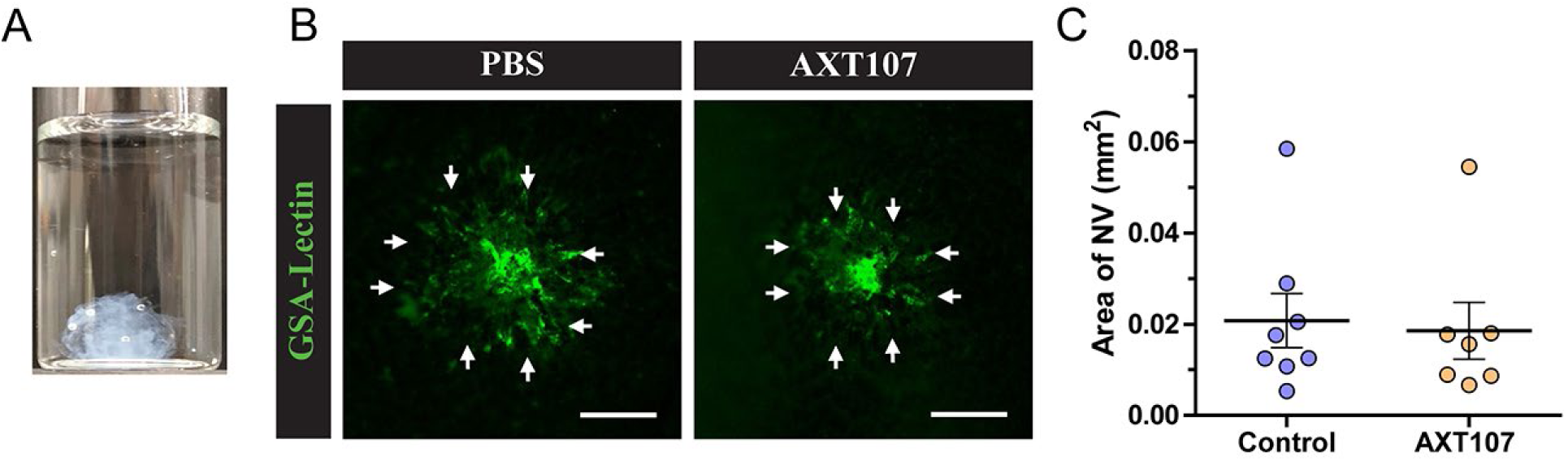
Suprachoroidal administration of soluble AXT107 does not inhibit CNV in a rat model. A) Gel-like structure of 500 µg soluble AXT107 injected into a vitreous substitute. B) Representative images of isolectin stained region of photocoagulation-induced NV in rat eyes treated by suprachoroidal injection with either PBS (left) or AXT107 (right). C) Quantification of the NV area from eyes described in (B). Scale bars = 100 µm. The areas of four target spots were averaged per eye and plotted; N= 7 or 8 (one mis-injection of an AXT107 eye). Differences were nonsignificant by Mann-Whitney test.

### Development and characterization of AXT107 microparticles

To improve the release of AXT107 from the suprachoroidal space and enhance its accumulation in retinal tissues, AXT107 was reformulated into a microparticle formulation (MP-AXT107). We hypothesized that increasing the surface area of the formulation, relative to the single gel-like mass formed by soluble AXT107, would promote hydration and facilitate release of AXT107 molecules.

MP-AXT107 was generated by exploiting the peptide’s limited solubility in ionic solutions. AXT107 dissolved in water was combined with a solution containing sucrose and NaCl under constant stirring, which was maintained overnight. Whereas soluble AXT107 exhibited an opalescent appearance, MP-AXT107 formed an opaque, white suspension (Fig. 2A). To quantify incorporation efficiency, particles were pelleted by centrifugation, and the supernatant was analyzed for soluble peptide content. Relative to a standard curve generated from equivalent amounts of soluble AXT107, only 0.59 ± 0.02% (mean ± SD) of peptide remained in the supernatant, indicating near-complete incorporation into the particulate fraction.

**Figure 2.**
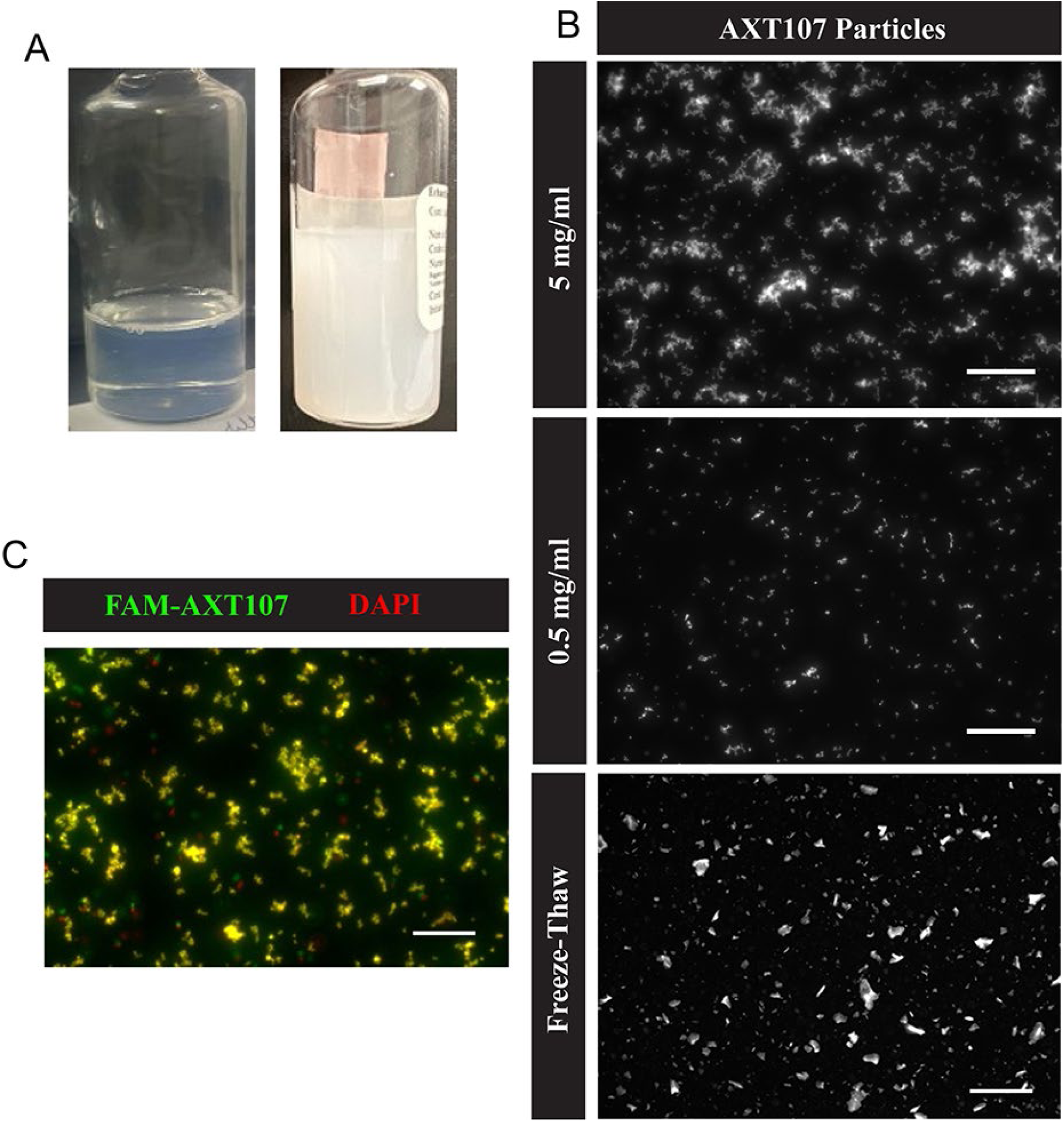
Characterization of MP-AXT107. A) Images of AXT107 solution (left) and MP-AXT107 (Right). B) Images of DAPI-stained MP-AXT107 at 5 mg/ml (top), 0.5 mg/ml (middle), and diluted to 0.25 mg/ml after 1 round of freeze-thaw (bottom). C) DAPI-stained (red) images of 0.5 mg/ml MP-AXT107 with 5% of AXT107 substituted for FAM-labeled AXT107 (green) with areas of overlap shown in yellow. Scale bars = 250 µm.

Because the opalescent appearance of soluble AXT107 suggests a supersaturated state, we evaluated whether centrifugation alone could induce peptide loss independent of particle formation. Surface samples of soluble AXT107 were collected before and after centrifugation and normalized to the mean pre-centrifugation concentration (set to 100%). The pre- and post-centrifugation samples measured 100 ± 12.1% and 100.6 ± 17.2% (mean ± SD), respectively, indicating no appreciable change in concentration. These results support the notion that the reduction in supernatant peptide observed for MP-AXT107 was specifically due to incorporation into particles.

Particle morphology was visualized by fluorescence microscopy following staining with DAPI, which was observed, but unreported, in previous work to fluoresce upon binding to self-assembled, but not soluble, AXT107 ^22^. At concentrations ≥5 mg/mL, the precipitated peptide formed loosely flocculated aggregates which transitioned to more uniform particles upon dilution (Fig. 2B). Early particle size measurements obtained by image analysis indicated an average diameter of 5.95 ± 2.97 µm (mean ± SD). Freezing the particles resulted in an irreversible increase in average diameter to 21.52 µm.

To confirm the specificity of the DAPI staining, particles were formulated with 5% 5-carboxyfluorescein (FAM)-labeled AXT107 (FAM-AXT107), which demonstrated near-complete colocalization between DAPI and FAM signals except for particles that shifted position during filter changes (Fig. 2C).

### AXT107 Particle Formulations Exhibit Long-term Stability

Long-term assessments of particle size and AXT107 chemical stability were conducted using GMP-quality MP-AXT107 drug product (Table 1). MP-AXT107 was prepared at concentrations of 1.25, 2.5, and 5 mg/mL AXT107 and stored in climate-controlled stability chambers at 5 ± 3 °C or 25 ± 2 °C/60 ± 5% relative humidity. Samples were collected at the indicated time points for analysis. Chemical stability was evaluated by quantifying full-length AXT107 and detecting impurities using HPLC relative to an AXT107 reference standard.

**Table 1.** Chemical and particle size stability of GMP MP-AXT107.

| Intended Conc. | Storage Temp. | Specification | Acceptable Range | Initial | 1 month | 3 months | 6 months | 9 months |
| --- | --- | --- | --- | --- | --- | --- | --- | --- |
| 1.25 mg/ml | 2-8 °C | Conc. (mg/ml) | 1.06-1.44 | 1.23 | 1.2 | 1.21 | 1.19 | 1.19 |
|  |  | PSA - D50 (µm) | n/a | 2.004 | 2.027 | 1.982 | 2.004 | 2.073 |
|  | 25 °C | Conc. (mg/ml) | 1.06-1.44 | 1.23 | -- | 1.19 | -- | -- |
|  |  | PSA - D50 (µm) | n/a | 2.004 | -- | 2.268 | -- | -- |
| 2.5 mg/ml | 2-8 °C | Conc. (mg/ml) | 2.13-2.88 | 2.26 | 2.13 | 2.13 | 2.35 | 2.26 |
|  |  | PSA - D50 (µm) | n/a | 1.791 | 1.895 | 1.832 | 1.853 | 1.895 |
|  | 25 °C | Conc. (mg/ml) | 2.13-2.88 | 2.26 | -- | 1.95* | -- | -- |
|  |  | PSA - D50 (µm) | n/a | 1.791 | -- | 1.916 | -- | -- |
| 5 mg/ml | 2-8 °C | Conc. (mg/ml) | 4.25-5.75 | 4.89 | 4.46 | 4.42 | 5 | 4.9 |
|  |  | PSA - D50 (µm) | n/a | 1.811 | 1.853 | 1.811 | 1.853 | 1.895 |
|  | 25 °C | Conc. (mg/ml) | 4.25-5.75 | 4.89 | -- | 3.78* | -- | -- |
|  |  | PSA - D50 (µm) | n/a | 1.811 | -- | 2.004 | -- | -- |

\* Value outside of acceptable range (±15% of target)

By 3 months, all samples stored at 25 °C exhibited measurable loss of AXT107, with greater percent loss observed at higher starting concentrations. Among the tested conditions, only the 1.25 mg/mL samples remained within the predefined specification of ±15% of the target concentration. In contrast, samples stored at 2–8 °C demonstrated improved stability, with all concentrations remaining within specification throughout the 9-month study period. Although some variability in measured concentrations was observed over time, no consistent downward trend was evident, suggesting that fluctuations were more likely attributable to assay variability rather than progressive peptide degradation.

Particle size stability was assessed using laser diffraction. Mean particle diameters ranged from 1.79 to 2.27 µm and remained consistent across all time points for samples stored at 2–8 °C. Samples stored at 25 °C exhibited greater variability in particle size over time; however, no clear directional trend was observed. Collectively, these data indicate that the MP-AXT107 remains physically and chemically stable for up to 9 months when stored at 2–8 °C.

### Efficacy of suprachoroidal AXT107 microparticles

The efficacy of MP-AXT107 was assessed using the same rat laser-induced CNV model as described above for the AXT107 solution. In this study rats received 3 μl injections of either vehicle (5% sucrose and saline) or 30 μg MP-AXT107. GSA-lectin staining of retinal flat mounts clearly show a reduction of CNV area in rats treated with AXT107 (Fig. 3A). Quantification of the total CNV per eye subsequently confirmed a significant, 60% reduction of neovascularization with MP-AXT107 treatment (Fig. 3B).

**Figure 3.**
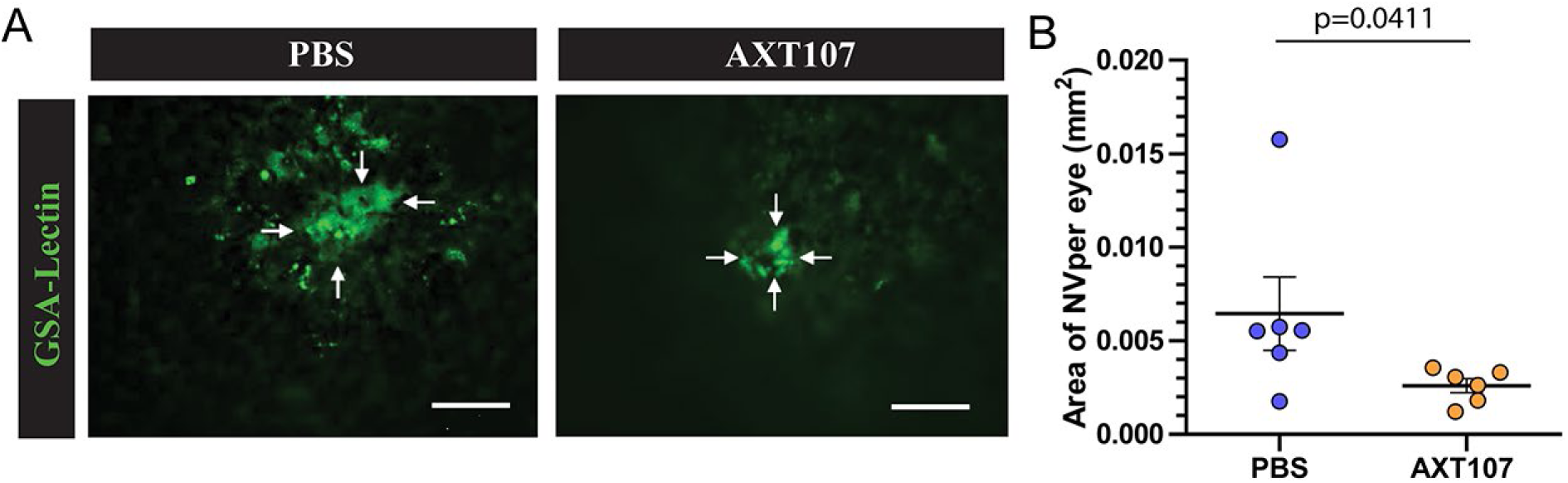
Suprachoroidal administration of MP-AXT107 significantly inhibits CNV in a rat model. A) Representative images of isolectin stained region of photocoagulation-induced NV in rat eyes treated by suprachoroidal injection with either PBS (left) or MP-AXT107 (right). C) Quantification of the NV area from eyes described in (A). Scale bars = 100 µm. The areas of four target spots were averaged per eye and plotted; N=6; p=0.0411 by Mann-Whitney test.

### GLP safety assessment of AXT107 MPs in a minipig single-dose toxicology study

Preclinical safety of suprachoroidal MP-AXT107 was evaluated in minipigs under GLP-compliant toxicology conditions over a 9-month study period. Given the absence of significant adverse findings across the full, extensive dataset, only the most relevant results to formulation safety are presented here. Animals received a single 100 µL suprachoroidal injection of vehicle (5% sucrose and saline) or MP-AXT107 at doses of 0.125 mg, 0.5 mg, or 1.25 mg. There was a noticeable increase in injection resistance with increasing dose, suggesting elevated viscosity, but did not require modification of the administration procedure. Successful delivery to the suprachoroidal space was confirmed by OCT imaging, which demonstrated a dose-dependent expansion of the suprachoroidal space (Fig. 4A), which is consistent with previously reported effects of viscosity on SCS injections.

**Figure 4.**
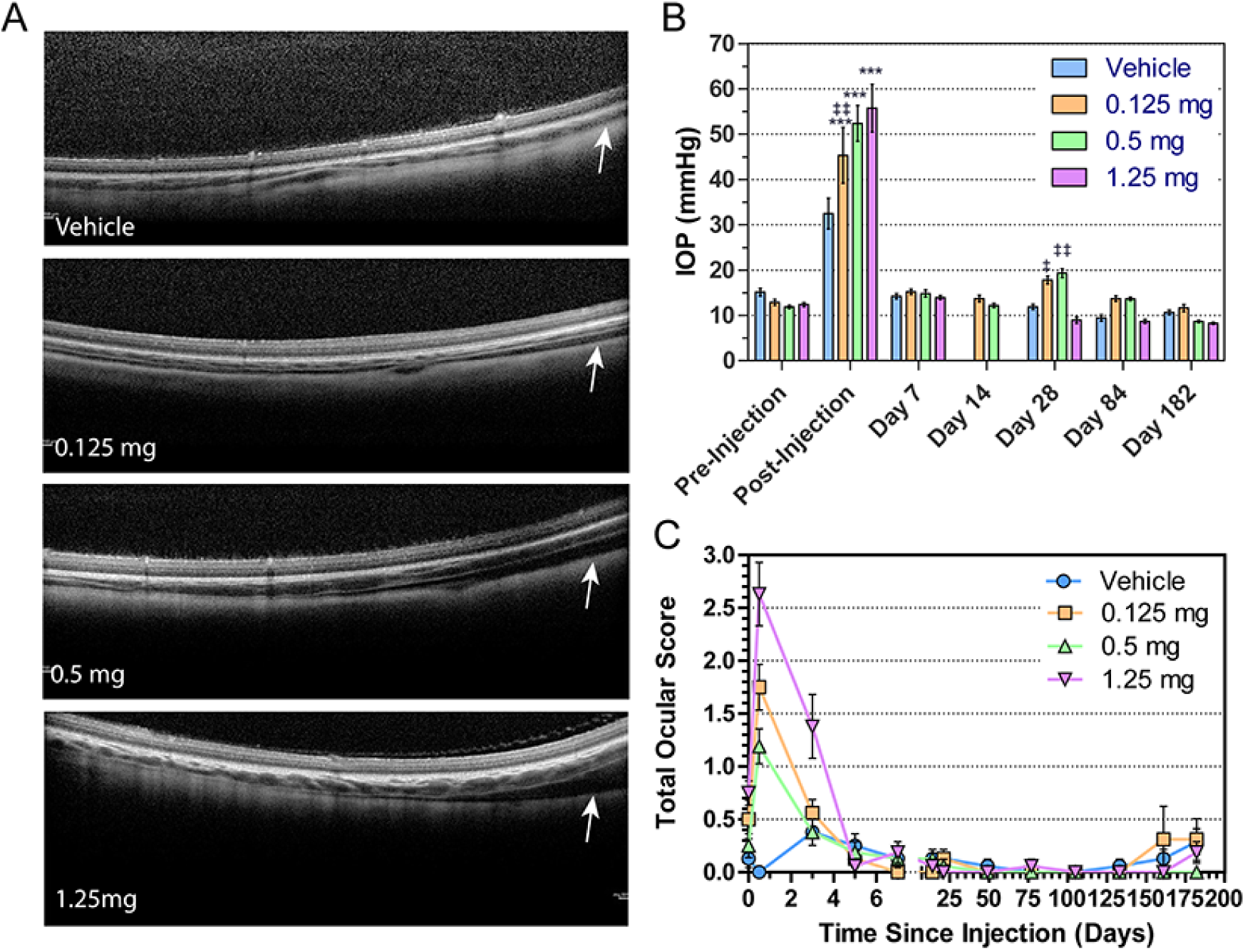
Single-dose toxicology study of MP-AXT107 in minipigs. A) OCT images of eyes following 100 µl suprachoroidal injections of vehicle (5% sucrose and saline) or varying doses MP-AXT107. White arrows indicated location of expanding suprachoroidal space. B) Average IOP per eye at different time points. Due to a scheduling error, D14 measurements are unavailable for vehicle and 1.25 mg. N=16 (8 animals); ***p<0.001 vs. vehicle; ‡ and ‡‡p>0.05 or 0.01 respectively vs. 1.25 mg AXT107 treatment. C) Average total ocular score per eye over time. N=16 (8 animals).

IOP exhibited an immediate, dose-dependent increase following injection (Fig. 4B), a transient effect commonly associated with suprachoroidal administration. IOP returned to baseline by the first post-dose assessment at one week and remained within normal limits for the duration of the study.

Ocular health was assessed longitudinally via slit-lamp examination by a qualified veterinarian and scored using a modified Draize method, incorporating evaluation of the cornea, iris, conjunctiva, lens, aqueous humor, and vitreous humor, with higher numbers reflecting greater severity (full criteria in Supplemental Table 1). Individual category scores were summed to generate a total ocular score per animal (out a maximal possible score of 32), and group mean scores were plotted over time (Fig. 4C). Treatment-related findings were primarily confined to the first few days post-injection and consisted predominantly of chemosis, corneal redness, and vitreous haze. These effects showed limited dose dependence and resolved to baseline for all groups by Day 5. Severity remained minimal even throughout these initial time points, with no animal exceeding a total ocular score of 4. Beyond this acute phase, ocular scores were consistently 0, with occasional scores of 1, for the remainder of the study. A single animal in the 0.125 mg group exhibited corneal findings beginning at Week 23 that persisted through the end of the study (Week 26). Given that these observations were limited to one animal in the lowest-dose group, these findings were considered independent of the treatment.

At study termination, the left eye from each animal was collected for bioanalysis. Eyes were dissected into whole tissue samples of aqueous humor, vitreous humor, retina, and choroid/RPE as well as sclera punches from 8 different sites, and AXT107 concentrations were quantified by mass spectrometry. AXT107 was below the limit of detection (<0.1 ng/g tissue) for all aqueous humor, retina, and vitreous humor samples, except for one retina sample from the 1.25 dose that contained 3523 ng AXT107/g tissue. Conversely, AXT107 was detectable in all but one of the choroid/RPE samples (ignoring one sample from each dose that were lost during preparation) and 27% of scleral samples. The mean value for each tissue sample per treatment group is shown in Table 2. As concentrations below the limit of detection or lost during processing were not included in the mean calculations, the total number of values included is also indicated. The levels of AXT107 in the choroid/RPE samples were similar among all treatment groups. The most consistent trend in the scleral samples appeared to be the detection of AXT107 in the superior and temporal regions, which coincides with the site of administration. AXT107 was only completely undetectable in one treated animal, from the 0.125 mg group, at the end of the 6-month study.

**Table 2.** Bioanalysis of minipig choroid and scleral tissues 9-months after MP-AXT107 Injection.

| *Tissue* |  | **Choroid/RPE** | **Sclera** |  |  |  |  |  |  |  |
| --- | --- | --- | --- | --- | --- | --- | --- | --- | --- | --- |
| *Region\** |  | *All* | *Ant. Inf. (1)* | *Ant. Inf. (2)* | *Ant. Sup. (1)* | *Ant. Sup (2)* | *Sup. Nasal* | *Inf. Nasal* | *Sup. Temp.* | *Inf. Temp.* |
| Vehicle | # Values | 0 | 0 | 0 | 0 | 0 | 0 | 0 | 0 | 0 |
|  | Mean (ng/g) | BQL | BQL | BQL | BQL | BQL | BQL | BQL | BQL | BQL |
|  | Std. Deviation | BQL | BQL | BQL | BQL | BQL | BQL | BQL | BQL | BQL |
| 0.125 mg | # Values | 6 | 1 | 0 | 3 | 0 | 6 | 4 | 5 | 0 |
|  | Mean (ng/g) | 647.2 | 94.6 | BQL | 88.83 | BQL | 79.42 | 110.9 | 150 | BQL |
|  | Std. Deviation | 377 | 0 | BQL | 44.6 | BQL | 37.61 | 107.4 | 102.6 | BQL |
| 0.5 mg | # Values | 7 | 1 | 0 | 2 | 1 | 3 | 0 | 5 | 0 |
|  | Mean (ng/g) | 531.4 | 94.7 | BQL | 88.7 | 98.7 | 118.2 | BQL | 112.8 | BQL |
|  | Std. Deviation | 375.4 | 0 | BQL | 52.75 | 0 | 131.8 | BQL | 57.33 | BQL |
| 1.25 mg | # Values | 7 | 2 | 1 | 2 | 4 | 1 | 1 | 5 | 0 |
|  | Mean (ng/g) | 653.6 | 42 | 152 | 3155 | 1376 | 76.7 | 131 | 109.6 | BQL |
|  | Std. Deviation | 735.8 | 8.344 | 0 | 4191 | 2475 | 0 | 0 | 75.84 | BQL |

**Abbreviations:** Ant. - Anterior; BQL - Below quantitation limit (0.1 ng/g tissue); Inf. - Inferior; Sup. - Superior; Temp. – Temporal
\* Choroid/RPE was processed as a whole while the sclera was processed as punches from 8 different regions. Two punches were made per anterior region and analyzed separately (labeled 1 and 2 above)

Systemic exposure was assessed via blood sampling throughout the study and subsequent bioanalysis. AXT107 concentrations were universally below the limit of detection (5 ng/mL), supporting minimal systemic exposure.

## Discussion

Anti-VEGF therapies are highly effective for retinal vascular diseases but are limited by the need for frequent dosing. Prior sustained-delivery approaches have been associated with anterior segment exposure and increases in IOP (NCT04697758, NCT04746963, and NCT03953079). To address these limitations, we evaluated suprachoroidal space (SCS) administration as a strategy to localize drug delivery while enabling sustained release.

Our results demonstrate that MP-AXT107 delivered to the SCS effectively suppresses VEGF-mediated vascular leakage in a rat model. In contrast, soluble AXT107 administered via the same route did not produce a measurable therapeutic effect despite clear efficacy in intravitreal models. This difference is likely driven by anatomical and transport barriers associated with SCS delivery. Intravitreal administration allows AXT107 to form a depot in proximity to the retina and target vasculature, whereas drug delivered to the SCS must first traverse the choroid. Injectate distribution may further contribute as intravitreal delivery permits some control over depot localization, while SCS injections deposit material near the pars plana and rely on injection-driven forces and diffusion for posterior distribution.

Although intraocular convection supports this movement, transport in the SCS is known to be influenced by physicochemical properties such as viscosity^30^. In this context, depot formation by soluble AXT107 may both limit diffusion towards the posterior pole of the eye, reducing access to macula, as well as prevent sufficient release of AXT107 to traverse the choroid and reach the retina in efficacious amounts. Reformulation into a particulate form appears to overcome these barriers by increasing surface area to enhance drug release and reducing self-assembly, thereby improving distribution.

The particle fabrication process was designed to be robust and scalable, enabling reproducible production of stable formulations. This process has been successfully transferred to a commercial manufacturing organization for clinical drug product manufacture. The resulting microparticles exhibit consistent size and remain stable for at least nine months under refrigerated storage, with no significant changes in peptide integrity or particle size. Although some settling was observed, particularly at lower concentrations, it was readily reversible with simple end-over-end mixing, allowing uniform resuspension and consistent dosing.

Bioanalysis of the eyes at the end of the study revealed that AXT107 distribution was concentrated within choroid/RPE and scleral fractions. This was not unexpected as the suprachoroidal space is located between these two tissue layers. Importantly, nearly all the treated animals contained some detectable amount of AXT107 in these tissues after 9 months, emphasizing the potential for a durable response from a single administration of the microparticle formulation. Moreover, the one animal without any detectable levels of AXT107 is noted as having a partly subconjunctival injection with no apparent expansion of the SCS in OCT, suggesting that this lack of detectable peptide could be at least partially explained by a mis-injection.

Suprachoroidal administration of AXT107 microparticles demonstrated a favorable safety profile. A transient increase in IOP was observed immediately following injection but resolved within one week without lasting effects. Such transient elevations are commonly reported following SCS administration and often resolve within hours^31^; however, earlier post-injection measurements were not captured in this study, representing a limitation.

Ocular inflammation followed a similar pattern, with mild findings observed shortly after dosing that resolved within one week and remained stable thereafter. Based on these findings, no safety concerns were identified, and 1.25 mg, the highest dose tested, was established as the no-observed-adverse-effect level (NOAEL).

## Materials and Methods

### Reagents

AXT107 was manufactured at Polypeptide Laboratories Inc. (Torrence, CA) by solid-state synthesis and lyophilized.

### AXT107 Gelation Assay

To demonstrate the gelling properties of AXT107, 50 µl of 10 mg/ml AXT107 in 5% sucrose was injected into a glass shell vial containing a vitreous substitute (3.97 mM glucose, 2.97 mM lactate, and 0.2 mg/ml hyaluronic acid from bovine vitreous in phosphate buffered saline).

### Generation of AXT107 Microparticles

AXT107 MPs (containing 1.25 to 10 mg/ml AXT107 and 5% sucrose) were generated by precipitating an AXT107 solution with a salt solution under constant stirring. For the AXT107 solution (2x final concentration AXT107, 10% sucrose, and HCl (see below)), powdered peptide (including FAM-labeled where relevant) was dissolved as much as possible in a solution of sucrose by vortexing. This mixture was then acidified with HCl and vortexed again until no further clarification was observed. Finally, the solution was homogenized using a D1000 handheld homogenizer between settings 3 and 4 for 30 seconds and immediately filtered through a hydrophobic PVDF syringe filter. For particle formation, the AXT107 solution was precipitated at a 1:1 volume ratio with a solution of NaCl and NaOH equimolar to the HCl added under constant stirring by magnetic stir bar. Stirring was then continued overnight. The final product was stored at 4 °C until used.

### Animals

Norway Brown Rats (6-8 weeks old; equal male and female) were purchased from Charles River Laboratories (Wilmington, MA). All studies were performed in accordance with the Association for Research in Vision and Ophthalmology Guidelines for research animal use and approved by the IACUC of Johns Hopkins University.

Gottingen Minipigs (3-5 months old) were purchased from Ellegaard Gottingen Minipigs A/S (Dalmose, Denmark). All experiments were performed at Specific Pig S.L (Barcelona, Spain) under GLP compliance. Studies were approved by the Ethic Experimental Committee of Specific Pig and authorized by the Departament de Territori i Sostenibilitat la Generalitat de Catalunya (Department of Territory and Sustainability of the Government of Catalonia).

### Rat Suprachoroidal Injection Model

Choroidal neovascularization (CNV) was generated by photocoagulation from laser-induced rupture of Bruch’ s membrane. Briefly, nine 6-8 week-old Norway brown rats were anesthetized, pupils were dilated with 1% tropicamide (Alcon Laboratories, Inc.) and Bruch’s membrane was ruptured at the 3, 9, and 12 o’clock positions of the posterior pole with a 562-nm diode laser (75-µm spot size, 0.1 s duration, 120 mW) using the slit lamp delivery system of an OcuLight GL Photocoagulator (Iridex) and a handheld cover slide as a contact lens.

For both soluble AXT107 and microparticle studies, rats were split into two groups of 3 animals that each received a suprachoroidal injection of 3 µl into both eyes. Group 1 received injections of either 33 mg/ml AXT107 or MP-AXT107 (10 mg/ml AXT107) and Group 2 received control injections of PBS in both studies. For injections, rats were anesthetized with ketamine/xylazine, and the eyes visualized with a Zeiss stereo dissecting microscope. A 30-gauge needle on a 1 ml syringe was then inserted four-fifths through the sclera 1 mm from the posterior limbus to create a circumferential opening. A 34-gauge Hamilton syringe was then inserted into this opening and through the remaining sclera into the suprachoroidal space. The plunger was then depressed until the entire 3 µl sample was injected. The needle was then withdrawn, and a cotton-tipped applicator applied to the site for 30 seconds to limit sample reflux. Fundus imaging was then used to inspect the success of the injection. The eye was then treated with antibiotic ointment and the rat returned to their cage.

Fourteen days after injections, the 9 rats were euthanized to measure the area of CNV. Retinas were dissected, stained with fluorescein (FITC)-conjugated Griffonia simplicifolia isolectin B4 (GSA) (Vector Laboratories) for 45 min to detect neovascularization, and flat-mounted. Flat mounts were examined by fluorescence microscopy, and the area of each choroidal NV was measured by image analysis with Image-Pro Plus software (Media Cybernetics) by an observer masked with respect to experimental groups. For statistical analysis, values of choroidal NV were averaged for each eye before comparison.

### Quantification of AXT107 Incorporation into Microparticles

MP-AXT107 were made fresh as described above and diluted in water as needed for quantification). A sample of the diluted stock was taken for measurement while the remainder was centrifuged for 5 min at 14000 x g and the supernatant sampled for measurement. As a control, the AXT107 solution prior to particle formation was also diluted in water to the same volume and sampled before and after centrifugation. Peptide content was quantified using the Pierce Quantitative Fluorometric Peptide Assay (ThermoFisher) using dilutions of AXT107 solution as a standard curve.

### Imaging AXT107 Microparticles

AXT107 microparticles were diluted as needed in 5% sucrose and NaCl and stained by adding 1.5 µg/ml DAPI. If used, FAM-labeled AXT107 was incorporated during particle generation. Samples were then loaded into hemocytometers and imaged using an Olympus Ix70 microscope at 100x total magnification.

### Image Based Particle Size Analysis

Samples were diluted at least 10-fold and stained with DAPI as described above. Using ImageJ software (NIH), background was subtracted, images converted to binary, corrected with watershed processing, and their area determined through the Analyze Particle feature. Particles were assumed roughly circular in shape and the diameter calculated from the area measurements.

### Particle size analysis and Stability Measurements of GMP AXT107

Particle size and chemical stability measurements for the GMP AXT107 suspension were performed by KABS Laboratories Inc. (St. Hubert, Canada). Particle size was measured with an AccuSizer780. As needed samples were diluted in vehicle (5% sucrose and saline) to maintain particle concentrations within the coincidence level of the sensor between 2000 particles/ml and 12000 particles/ml.

For chemical stability, sealed sample vials containing AXT107 suspensions were stored in climate-controlled chambers at 5±3 °C or 25±2 °C/60±5% for the indicated amounts of time. At each timepoint, a sample vial was removed and diluted to 0.1 mg/ml in water/glacial acetic acid (20/80 v/v) to dissolve the particles. A reference standard was also generated by dissolving AXT107 powder in the water/glacial acetic acid mix to 0.1 mg/ml.

The amount of full length AXT107 and impurities were then assessed using Agilent 1100 or 1200 HPLC systems with UV detectors and Xbridge® C18 5μm, 4.6×150mm columns. The column was pre-equilibrated with 90:10 mix of mobile phase A (20mM NH4HCO3 in H2O/ACN: 90/10 (v/v)) and mobile phase B (20mM NH4HCO3 in H2O/ACN: 20/80 (v/v)) and the peptide eluted by a gradient up to 100% mobile phase B. The percentage of full-length AXT107 to impurities was determined by measuring the area of analyte peaks on the resulting UV chromatograms.

### Gottingen Minipig Preparation and Injection

Fourteen days before the study, Gottingen minipigs (3-5 months old) were examined by qualified study personnel and external veterinary ophthalmologists to ensure suitability and establish baseline values for monitored biological variables. Notably, these included IOP measurements and ocular health scoring (described in detail below) in addition to other ocular tests (not included here) and general health examinations. Thirty-two animals were then randomized into 4 study groups of 8 animals each according to body weight and sex (4 males and 4 females). Groups included vehicle control (5% sucrose and saline) and particle formulations containing 1.25 mg/ml, 5 mg/ml, or 12.5 mg/ml AXT107.

For treatment administration, animals were sedated with an intramuscular injection of 0.02 mg/kg dexmedetomidine, 0.3 mg/kg midazolam, and 0.3 mg/kg butorphanol and eyes treated with 0.5% tetracaine and 1% tropicamide for further anesthetization and pupil dilation. The conjunctival fornixes were flushed with a 1:50 dilution of iodine in saline solution and the eyelid margins swabbed with undiluted 5% betadine solution. Treatments were delivered to the suprachoroidal space using Bella-Vue 900 silicon microneedles (Uneedle, Enschede, Netherlands). Microneedles were inserted into the superotemporal quadrant 4 mm posterior from the limbus, and 100 µl of the test article was slowly administered over 5-10 seconds. A cotton-tipped applicator was then applied to the injection site for another 30 seconds to reduce reflux. This process was repeated for both eyes for final doses of 0 (vehicle), 0.125, 0.5, and 1.25 mg/eye.

### Minipig Intraocular Pressure measurements

Intraocular pressure was measured using a rebound tonometer (Tonovet-Plus) while the animal was under sedation as described for injections above. Interpretations of the results were performed by board-certified veterinary ophthalmologists. IOP was measured 2 weeks and immediately before article administration, immediately following, and after 1, 2, 4, 12, and 26 weeks.

### Ocular Health Scoring

The health of both eyes was scored using a modified Draize technique. The evaluation was performed by board-certified veterinary ophthalmologists using a slit lamp two weeks and immediately before test article administration, days 1, 3, 5, and 7 and weeks 2, 3, 7, 11, 15, 19, 23, and 26. A full list of the criteria and their corresponding values can be found in Supplemental Table 1.

### Toxicokinetics

Blood samples (2 ml) were collected from the jugular or cava veins into prechilled blood tubes containing EDTA K2 immediately before test article administration and 1, 4, 8, 12, 16, 20, and 26 weeks after. Samples were maintained on ice during processing, which included centrifugation and isolation of plasma. The plasma was mixed 1:1 in a solution of 0.8M ammonium formate and 2% CHAPS in reverse osmosis water and stored at -80 °C. Samples were analyzed within 1 month of collection.

AXT107 concentration is analyzed by an LC-MS/MS method at Origin Bioanalytical Laboratory, Inc. (Rancho Cordovo, CA). Plasma samples were treated with an appropriate volume of solvent 50:50 ethanol:water with 0.5% ammonium hydroxide. AXT107 and internal standard AXT107 F^+10 were extracted from 100 μL of the plasma sample by protein precipitation extraction. Calibration curves and analytical QCs were prepared in surrogate matrix [1:9 Minipig Plasma: (50:50 ethanol:water) w/ 0.5% NH4OH]. Process QCs in each matrix were homogenized and extracted alongside study samples. The final extracts were then analyzed using reverse phase LC-MS/MS.

### Ocular Tissue Bioanalysis

Whole eyes were collected during necropsy by Specific Pig S. L. and marked by suture at the 12:00 position. Following enucleation, all procedures were performed on ice. Excess tissue was trimmed and as much of the aqueous humor collected. The globe was then washed in saline, blotted dry, and flash frozen in liquid nitrogen. All tissues and samples were then stored at -80 °C prior to analysis.

Separation of the remaining ocular tissues was performed at Lifecorp Early Development Laboratories Inc. (Madison, WI). Scleral samples were taken using an 8 mm biopsy punch (target 300-500 mg each punch) from four approximately equal quadrants (corresponding to the superior-temporal, superior-nasal, inferior-temporal, and inferior-nasal regions). The anterior sclera was trimmed from the cornea and collected as superior and inferior samples. The frozen eye was then bisected and the frozen vitreous samples removed and combined. The retina was gently lifted from both halves of globe using filter paper and combined. Residual scleral tissue was then dissected away and the remaining tissue collected as the choroid-RPE.

AXT107 tissue concentrations were analyzed by an LC-MS/MS method at Origin Bioanalytical Laboratories. Anterior scleral samples were further subdivided prior to analysis. Aqueous humor samples were treated with an appropriate volume of 0.5M CHAPS solution to recover analyte that may be lost due to non-specific binding. AXT107 and internal standard AXT107 F^+10 were extracted from 50 μL of the treated aqueous humor sample by protein precipitation extraction. Calibration curves and analytical QCs were prepared in surrogate matrix [0.5% BSA with 15 mM CHAPS]. Choroid-RPE, retina, sclera, and vitreous humor samples were homogenized in lysis buffer solution and an appropriate volume of solvent [(70:30 ethanol:water) w/0.7% NH4OH]. AXT107 and internal standard AXT107 F^+10 were extracted from 100 μL of the ocular sample homogenate by protein precipitation extraction. Calibration curves and analytical QCs were prepared in surrogate matrix [1:9 Minipig Plasma: (50:50 ethanol:water) w/ 0.5% NH4OH]. Process QCs in each matrix were homogenized and extracted alongside study samples. The final extracts were then analyzed using reverse phase LC-MS/MS.

## Acknowledgements

This project was supported with internal funding by AsclepiX Therapeutics, Inc. and NIH grant R01EY028996-06.

## Author Contributions

ACM, TJR, JJG, ASP, and NBP conceptualized the project and determined experimental methods. ACM, TJR, and NBP developed and characterized the particle formulations, and monitored work at external facilities. RL, JS, and PAC developed the rat model and associated protocol; RL and JS performed rat studies. All authors assisted in analyzing data. ACM wrote the manuscript with input from the other authors.

---

**Supplemental Table 1.** Ocular scoring guidelines.

| Eye Strucure | VARIABLE | DESCRIPTION | SCORE |
| --- | --- | --- | --- |
| CORNEA | Opacity- degree of density | No opacity | 0 |
|  |  | Scattered or diffuse area, details of iris clearly visible | 1+ |
|  |  | Easily discernible translucent areas, details of iris slightly obscured | 2+ |
|  |  | Opalescent areas, no details of iris visible, size of pupil barely discernible | 3+ |
|  |  | Opaque, iris not visible | 4+ |
|  | Area of cornea involved | No opacity (no involvement) | 0 |
|  |  | One quarter (or less) but not zero | 1 |
|  |  | Greater than one quarter, but less than a half | 2 |
|  |  | Greater than one half, but less than three quarters | 3 |
|  |  | Greater than three quarters, up to whole area | 4 |
| IRIS | Values | Normal | 0 |
|  |  | Folds above normal, congestion, swelling, circumcorneal injection (any or all of these of combination of any thereof), iris still | 1+ |
|  |  | No reaction to light, haemorrhage, gross destruction (any or all of these) | 2+ |
| CONJUNCTIVA | Redness<br>(refers to palpebral and bulbar conjunctivae excluding cornea and iris) | Vessels normal | 0 |
|  |  | Vessels definitely injected above normal | 1 |
|  |  | More diffuse, deeper crimson red, individual vessels not easily discernible | 2+ |
|  |  | Diffuse beefy red | 3+ |
|  | Chemosis | No swelling | 0 |
|  |  | Any swelling above normal (includes nictitating membrane) | 1 |
|  |  | Obvious swelling with partial eversion of lids | 2+ |
|  |  | Swelling with lids about half closed | 3+ |
|  |  | Swelling with lids about half closed to completely closed | 4+ |
|  | Discharge | No discharge | 0 |
|  |  | Any amount different from normal (does not include small amounts observed in inner canthus of normal animals) | 1 |
|  |  | Discharge with moistening of the lids and hairs just adjacent to lids | 2 |
|  |  | Discharge with moistening of the lids and hairs, and considerable area around the eye | 3 |
| AC flare | Total number of AC cells viewed in volume (field) of the slit beam | No cells in field | 0 |
|  |  | 1-25 cells in field | 1+ |
|  |  | 26-50 cells in field | 2+ |
|  |  | 51-100 cells in field | 3+ |
|  |  | >100 cells in field | 4+ |
| LENS | Opacification/cellular infiltrate | Normal | 0 |
|  |  | Lens opacification/cellular infiltrate (incipient <5%)/cellular infiltrate | 1+ |
|  |  | Lens opacification/cellular infiltrate (early immature 5-50%) | 2+ |
|  |  | Lens opacification/cellular infiltrate (late immature 50-95%) | 3+ |
|  |  | Lens opacification/cellular infiltrate (complete) | 4+ |
| Vitreous haze | Vitreous haze | Normal | 0 |
|  |  | Mild vitreous haze | 1+ |
|  |  | Moderate vitreous haze | 2+ |
|  |  | Marked vitreous haze | 3+ |
|  |  | Severe vitreous haze | 4+ |

